# Stress-induced epigenetic regulation of transcription in neocortical excitatory neurons drives depression-like behavior

**DOI:** 10.1101/2020.07.06.190280

**Authors:** Deborah Y. Kwon, Peng Hu, Ying-Tao Zhao, Jonathan A. Beagan, Jonathan H. Nofziger, Yue Cui, Bing Xu, Daria Zaitseva, Jennifer E. Phillips-Cremins, Julie A. Blendy, Hao Wu, Zhaolan Zhou

## Abstract

Prolonged stress exposure is a major risk factor for the development of depression and comorbid anxiety. Efforts to understand how recurrent stress induces behavioral maladaptation have largely concentrated on the neuronal synapse with limited understanding of the underlying epigenetic mechanisms. Here, we performed complementary bulk nuclei- and single-nucleus transcriptome profiling and mapped fine-scale chromatin architecture in mice subjected to chronic unpredictable stress (CUS) to identify the cell type-specific epigenetic alterations that drive complex behavior. We find that neocortical excitatory neurons are particularly vulnerable to chronic stress and acquire signatures of transcription and chromatin configuration that denote reduced neuronal activity and expression of Yin Yang 1 (YY1). Selective ablation of YY1 in excitatory neurons of the prefrontal cortex (PFC) enhances stress sensitivity in mice, inducing depressive- and anxiety-related behaviors and deregulating the expression of stress-associated genes following an abbreviated stress exposure. Notably, we find that loss of YY1 in PFC excitatory neurons provokes more maladaptive behaviors in stressed females than males. These findings demonstrate how chronic stress provokes maladaptive behavior by epigenetically shaping excitatory neurons in the PFC, identifying a novel molecular target for therapeutic treatment of stress-related mood and anxiety disorders.

## INTRODUCTION

Major depressive disorder (MDD) represents the leading global cause of disability^1^. Its high comorbidity with mental disorders like anxiety and rising prevalence worldwide^1,2^ necessitate further understanding of its etiology. An abundance of epidemiological studies documenting MDD onset following adverse life experiences have led to its categorization as a stress-related illness^3–5^Notably, rodents exposed to various forms of chronic stress also exhibit depressive- and anxiety-related phenotypes^6,7^. Female rodents even show elevated endocrine and behavioral responses to stress compared to males^8,9^, consonant with the increased incidence of stress-related disorders in women^10–12^. These findings underscore an evolutionarily conserved effect of stress on the brain and behavior, providing a basis for modeling the pathophysiology of stress-related mood and anxiety disorders in laboratory animals.

The prefrontal cortex (PFC)—the brain region responsible for top-down regulation of cognition, emotion, and behavior—is linked to depression and acutely sensitive to trauma and stress exposure^13^. A concatenation of preclinical and clinical studies support PFC dysfunction in MDD and other anxiety-related disorders^14–17^, and a number of structural and functional changes in PFC pyramidal neurons—the primary glutamatergic excitatory cells in this brain region—have been reported in animals experiencing sustained stress or chronic glucocorticoid exposure^18–23^. Decreased grey matter volume^24–26^, synapse number^27^, and altered glutamate levels^28,29^ in the PFC have also been reported in MDD patients. These findings, taken together, have led to a glutamate hypothesis of depression, which theorizes that the disruption of glutamate-excitatory neurotransmission in the PFC leads to PFC hypoactivity, dysfunction and impaired emotional regulation.

While the functional impact of chronic stress on the excitatory synapse is well studied, its effect on the nucleus—where epigenetic processes that dynamically regulate transcription occur—that underlie or complement these synaptic alterations are poorly understood. Recent advancements in next-generation sequencing technology have enabled genome-wide transcriptional profiling in select brain regions of chronically stressed animals and MDD patients. These studies report broad alterations in whole cell RNA populations across several brain regions associated with chronic stress and disease^9,30–32^, suggesting that epigenetic changes underlie the manifestation of stress-related mood disorders. While these findings inform us of the general effects of chronic stress on steady-state RNA levels, they cannot distinguish between alterations made in the nucleus during transcription or by post-transcriptional mechanisms in the cytoplasm. Moreover, the brain is a heterogeneous organ composed of numerous discrete cell types that are defined by unique transcriptomes. Chronic stress likely impacts transcription in a cell type-specific manner that cannot be fully resolved by profiling RNAs isolated from bulk tissues.

To probe the epigenetic mechanisms in the PFC that drive behavioral maladaptation to chronic stress, we performed complementary bulk and single-cell sequencing of nuclear RNA transcripts and mapped activity-dependent changes in chromatin architecture. This multipronged approach reveals key transcriptional regulators and gene-regulatory networks in neocortical excitatory neurons altered by chronic stress exposure. We show that these effects serve to shape these cells into a state of hypoactivity by reducing the transcription of synaptic genes involved in glutamatergic neurotransmission and restructuring genome architecture into a pattern associated with synaptic inactivity. We find that these alterations are mediated, in part, by a corticosterone-induced reduction of the transcriptional regulator, Yin Yang 1 (YY1), leading to the transcriptional misregulation of other transcription factors and compounding the effects of chronic stress on these cells. Using adenoviral-mediated inactivation of *Yy1* in PFC excitatory neurons, we identify a novel cell type-specific role for YY1 as a critical regulator of stress coping in male and female mice. Together, these findings provide novel insight into the epigenetic mechanisms that transpire within the nuclei of neocortical excitatory neurons to drive maladaptive transcriptional and behavioral responses to chronic stress.

## RESULTS

### Twelve days of chronic unpredictable stress drives depressive- and anxiety-like behaviors in adult male mice

Considering the preponderance of epidemiological data linking chronic stress to major depressive and anxiety disorders, we first characterized the direct effects of chronic stress on behavior. We chose to use the chronic unpredictable stress (CUS) rodent model of depression due to its previously reported capacity to induce anhedonia and long-term depression-related phenotypes in adult male and female mice^33,34^. We expanded upon these findings by performing a battery of behavioral tests to assay depression- and anxiety-associated behaviors.

CUS consists of varying stressors delivered three times daily for variable lengths of time to prevent stress habituation (Supplemental Table 1). We subjected a cohort of adult male mice (9-10 weeks old) to twelve consecutive days of CUS followed by behavioral testing to assess the efficacy of the CUS paradigm. We first examined our mice for changes in body weight and food consumption, as stress exposure is known to affect these measures. Body weights of control and CUS animals were evenly distributed prior to CUS exposure (Fig 1a). However, we found that twelve consecutive days of CUS produced significant changes in body weight between the two age-matched cohorts (P<0.0001). While mice in the non-stressed control group gained ~1g of weight on average over the twelve-day period, CUS prevented weight gain and produced weight loss in ~67% of the CUS cohort (Fig 1b). We also found that CUS males consumed significantly less food than controls, even when food consumption was normalized to body weight to account for weight loss (P<0.01; Fig 1c).

**Figure 1.**
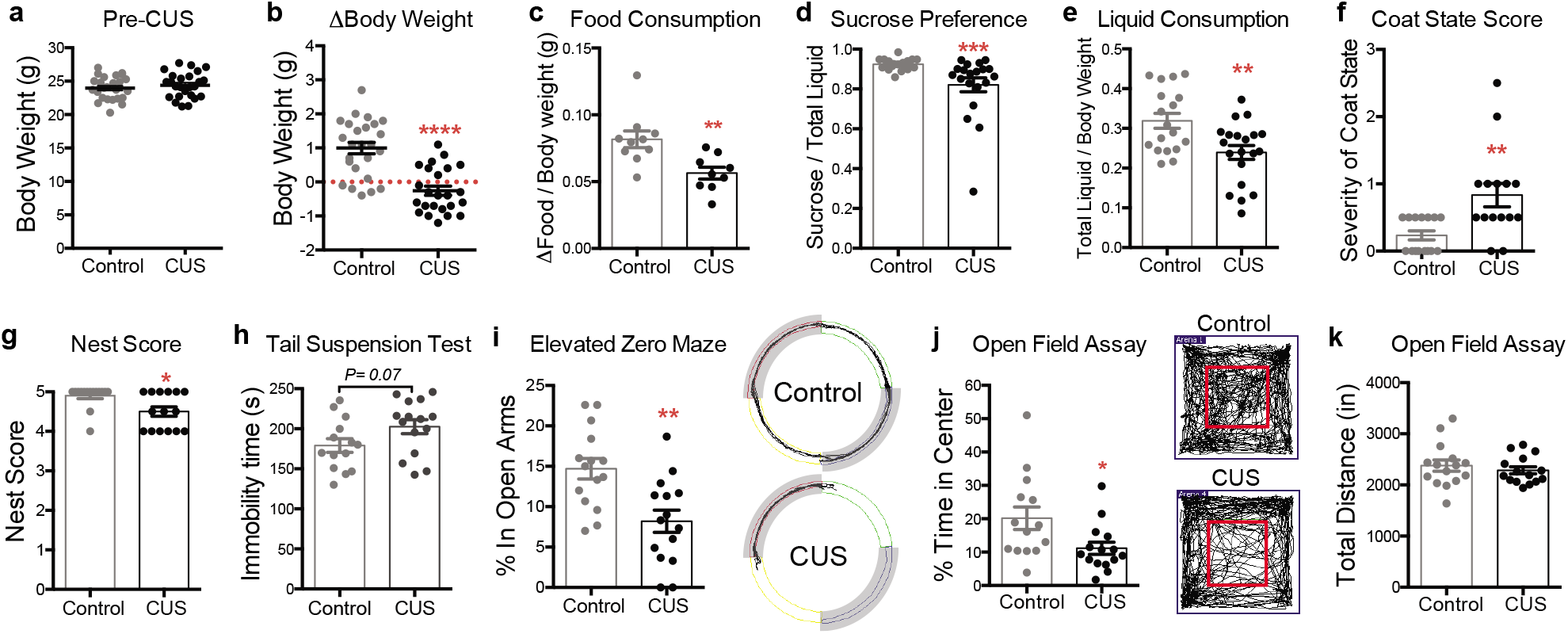
Twelve days of chronic unpredictable stress (CUS) drives a battery of depressive- and anxiety-like behaviors in adult male mice. (a) Pre-CUS body weights of 9-10 weeks old control and CUS males (n=25 per group). (b) CUS induces weight loss in adult males (Student’s t-test; P=0.0001; n=20 per group; error bars, s.e.m.). (c) Food consumption of control and CUS males normalized to body weight (Mann-Whitney U-test; P=0.0021; n=10 controls, n=9 CUS). (d) CUS decreases sucrose preference in male mice relative to controls (Mann-Whitney U-test; P=0.0004; n=18 controls, n=20 CUS). (e) Liquid consumption of control and CUS males normalized to body weight (Mann-Whitney U-test; P=0.004; n=18 controls, n=20 CUS). (f) CUS increases severity of coat scores relative to controls (Mann-Whitney U-test; P=0.002; n=15 per group). (g) CUS decreases nest scores relative to controls (Mann-Whitney U-test; P=0.01; n=15 per group). (h) CUS males show a trend towards enhanced immobility in the tail suspension test (Student’s t-test; P= 0.067; n=14 controls, n=15 CUS). (i) CUS males show decreased percent time spent in the open arms of the elevated zero maze (Student’s t-test; P= 0.0017; n=15 per group). Representative tracks obtained from video-tracking software are shown on the right. Closed arms are shaded in grey. (j) CUS males show decreased percent time spent in the center of the open field arena (Student’s t-test with Welch’s correction; P= 0.03; n=14 controls, n=15 CUS) without altered locomotor activity (P=0.5). Representative tracks obtained from video-tracking software are shown in the middle, with the boundaries of the center of the arena demarcated in red. *P<0.05; **P<0.01; ***P<0.001; ****P<0.0001. Error bars represent s.e.m.

We next assayed control and CUS mice for anhedonia, a hallmark symptom of depression, using the sucrose preference test. CUS males showed significantly decreased sucrose preference than controls (P< 0.001; 1d) and were also found to consume less total liquid over a 24-hour period (P< 0.05; 1e). Loss of motivation is also associated with depression. Accordingly, we evaluated coat state, an indirect measure of grooming behavior, and nest-building to assess motivated behaviors in CUS-subjected animals^35,36^. We found that CUS mice exhibited deteriorated coat states, as reflected by significantly higher coat state scores (P< 0.01; Fig 1f), as well as decreased nest scores after a 16-hour overnight period with a fresh, intact cotton nestlet (P< 0.05; Fig 1g). However, nests between control and CUS animals were virtually indistinguishable after 24 hours (data not shown), indicating that CUS induces a delay in nest-building and not physical impairment in nest-building ability. Lastly, behavioral despair, a depressive-like phenotype measured by the tail suspension test (TST), showed a trend towards increased immobility compared to controls (P= 0.07; Fig 1h).

Given the high comorbidity of anxiety and depression, we also assessed anxiety-related behaviors in CUS animals using the elevated zero maze (EZM) and the open field tests (OFT), both of which measure exploratory behavior related to anxiety. CUS mice were found to spend less time in the open arms of the elevated zero maze (P< 0.01; Fig 1i), as well as reduced time in the center of the open field arena in the OFT (P< 0.05; Fig 1j). Importantly, control and CUS mice traveled comparable distances in the OFT, indicating that the observed reduction in exploratory behavior was not due to CUS-induced differences in physical activity (Fig 1k). Taken together, these findings demonstrate that twelve days of CUS produces robust depressive- and anxiogenic-like phenotypes in adult male mice.

### CUS deregulates transcription and alters chromatin folding in the prefrontal cortex

We next sought to understand the transcriptional mechanisms underlying the observed effects of CUS on behavior. Gene expression changes can have a profound impact on neuronal physiology, and broad genomewide alterations in gene expression have been reported in MDD and chronically stressed animals^9,30,31^. However, the vast majority of these RNA profiling experiments have assayed whole cell RNA that provide an overview of gene expression changes but are unable to distinguish between those that arise during or posttranscription. Nuclear transcriptomes, which are mostly comprised of nascent pre-mRNA transcripts, provide a better representation of Pol II activity and chromatin state^35,36^. Thus, we performed RNA-sequencing (RNA-seq) on nuclei isolated from medial PFC tissues of control and CUS animals to determine how chronic stress impacts the transcriptional landscape of the cells that compose this stress-sensitive brain region.

We profiled nuclear RNA transcripts from 9 individual adult male mice (4 unstressed controls, 5 CUS). Our analysis uncovered 1,362 differentially expressed transcripts in CUS samples (FDR <0.05, Supplemental Table 2), 832 of which are downregulated and 530 upregulated (Fig 2a; Supplemental Table 2). We found that the large majority of differentially expressed transcripts are composed of protein-coding genes (86.3%), but also identified differentially expressed long noncoding RNAs, including long interspersed noncoding RNAs (lincRNAs, 7.1%), antisense transcripts (3.4%), and noncoding processed transcripts (1.5%), as well as small non-coding RNAs, which include microRNAs (1.3%) and small nucleolar RNAs (snoRNAs, 0.37), and 1 mitochondrial RNA (mtRNA) (Fig 2b).

**Figure 2.**
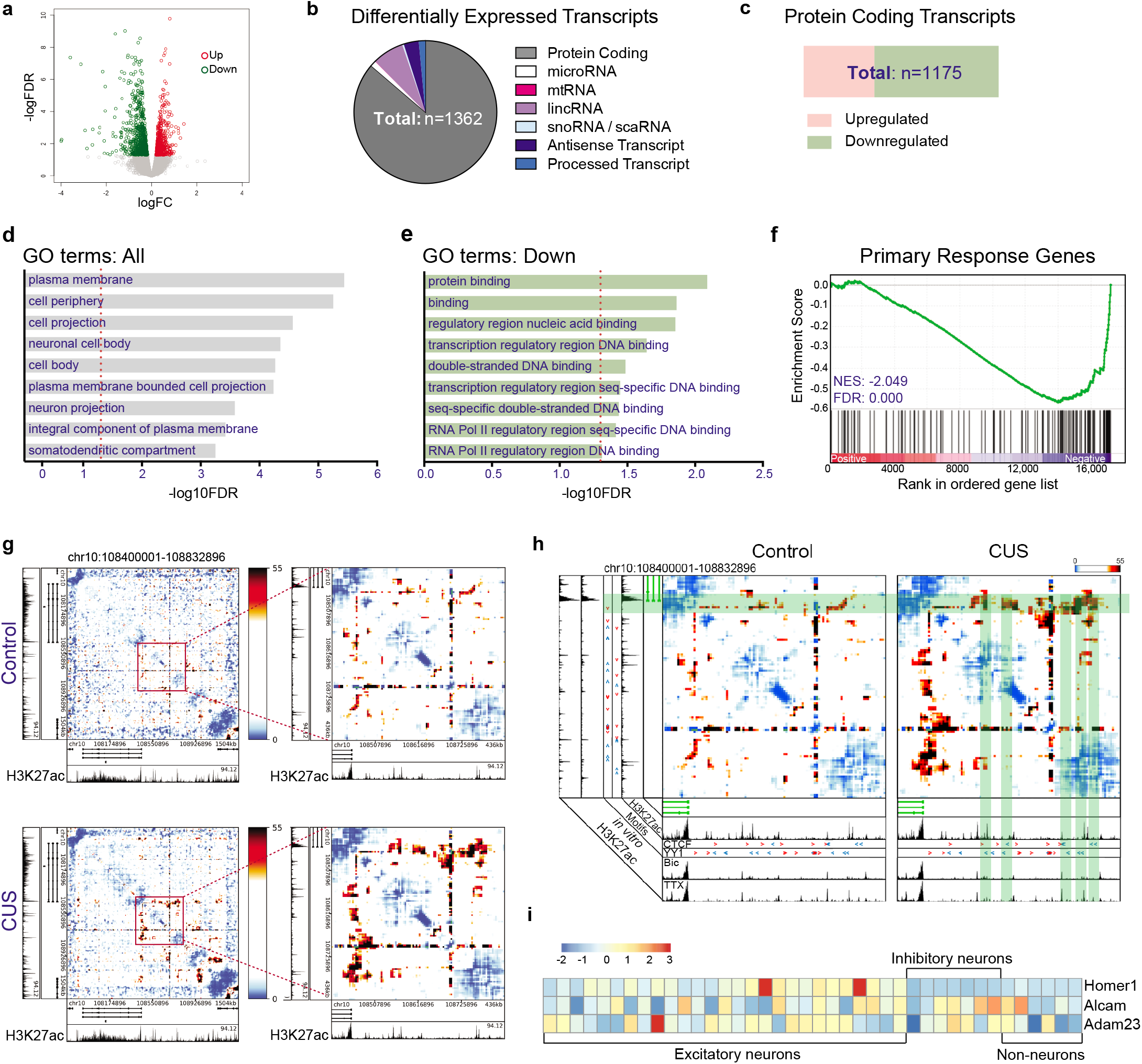
CUS impacts transcription and chromatin folding in the PFC. (a) Significantly upregulated transcripts (red) and downregulated transcripts (green) from comparison of control and CUS PFC nuclear transcripts (FDR<0.05). (b) Piechart exhibiting representation of differentially expressed RNA transcript populations. (c) Barchart portraying makeup of upregulated and downregulated differentially expressed protein-coding transcripts. (d,e) Top 9 most significantly enriched gene ontology (GO) terms for (d) all, and (e) down-regulated CUS DEGs (FDR<0.05). (f) GSEA plot charting negative enrichment of neuronal primary response genes in list of differentially expressed genes obtained from bulk nuclear RNA-seq analysis of control and CUS PFCs. NES, normalized enrichment score. (g) Background-corrected interaction frequency heatmaps displaying chromatin contacts in a 1.5-Mb region surrounding the *Syt1* gene in control and CUS frontal cortical tissues. The highlighted region in each heatmap marks the location of a zoomed-in plot. H3K27ac ChIP-seq track from adult cortical excitatory neurons control is shown below each heatmap. (h) Heatmaps plotting genomic locations and directionality of CTCF and YY1 binding sequences (red and blue arrowheads) in regions showing increased chromatin interactions in CUS mice (green shaded boxes). (i) Heatmap showing expression of CUS DEGs, *Homer1, Adam23*, and *Alcam*, across the cortical cell populations defined by single-nucleus RNA-seq in the adult mouse neocortex (Hu et al., 2017). Legend depicts z-score of normalized gene expression.

Gene ontology (GO) analysis of the differentially expressed protein-coding genes (DEGs) identified an enrichment of neuron synapse and axonal-related terms (Fig 2d), indicating that a significant number of CUS-associated DEGs are found in neurons. We also found that down-regulated genes, representing the majority of all CUS-associated DEGs (63.7%; Fig 2c), primarily encode proteins related to DNA binding and Pol II-mediated transcription (Fig 2e).

We observed that several of the down-regulated DNA binding genes that were identified by GO analysis, including *Fos, Fosb*, and *Fosl2*, are regulated by neuronal activity. These data suggests that activity-regulated gene expression programs in the PFC might be altered by CUS exposure. We subsequently performed a preranked gene-set enrichment analysis (GSEA^37^) to test for the overrepresentation of primary response genes, which are induced by neuronal stimulation^38^, in our RNA-seq data. Indeed, we found a negative enrichment of primary response genes (normalized enrichment score= −2.05; FDR= 0.000) in our analysis, indicating that CUS leads to decreased neuronal activity in the PFC (Fig 2f).

These findings led us to consider whether CUS induces a remodeling of higher-order genome architecture that reflect changes in neuronal activity. Genomes are organized into three-dimensional structures, forming chromatin fibers that can be dynamically looped by proteins such as CTCF, cohesin, and YY1 to bring together regulatory genomic sequences with their targets^39–41^. We leveraged Chromosome-Conformation-Capture-Carbon-Copy (5C) sequencing data generated from mouse cortical neurons that were pharmacologically modulated with treatments of bicuculline (Bic) or tetrodotoxin (TTX), which stimulate and inhibit neuronal activity, respectively, to map activity-dynamic chromatin loops. We observed chromatin interactions between the gene encoding the pre-synaptic membrane protein, *Synaptotagmin-1* (*Syt1*), and upstream activity-decommissioned enhancers associated with H3K27ac that increased in TTX-inactivated neurons and decreased in Bic-stimulated neurons relative to untreated cells (Supplemental Fig 1a). These data demonstrate that chromatin architecture and histone acetylation around the *Syt1* locus is dynamically remodeled by neuronal activity. Notably, we found that chromatin contacts at these same regulatory elements also increased in frontal cortices of CUS-subjected mice compared to controls, indicating that chronic stress exposure restructures genome organization at this locus into a pattern associated with TTX-inhibition of neuronal activity (Fig 2g, Supplementary Fig 1b). To identify the regulatory proteins accountable for these chromatin alterations, we also surveyed the genome upstream of *Syt1* for the presence of binding motifs of known architectural proteins—namely CTCF and YY1—which have known roles in organizing 3D chromatin structure. Using JASPAR, we uncovered YY1 and CTCF motif sequences in the regulatory regions showing increased chromatin interactions in CUS-subjected mice and TTX-treated cortical neurons (Fig 2h, red and blue arrowheads).

Taken together, these results indicate that CUS exposure dynamically shapes the PFC into a state of neuronal inactivity by decreasing the transcription of neuronal activity-dependent genes and restructuring higher-order genome architecture into a pattern associated with synaptic silencing. Furthermore, our findings suggest that the zinc-finger transcription factors, CTCF and YY1, mediate chromatin folding at these loci in response to chronic stress and TTX administration.

### Broad distribution of DEG expression in the neocortex

The cerebral cortex is a highly heterogeneous brain region composed of numerous neuronal and non-neuronal subtypes. Having determined that CUS alters transcription in cortical cells, we sought to construct cell typespecific models of transcriptional regulation by broadly classifying each DEG into a known cortical cell type. To accomplish this, we analyzed DEG expression across every cell type-specific cluster obtained in our previously published single-nucleus RNA-sequencing analysis of the mouse cortex^42^. However, we discovered that many CUS DEGs are expressed in multiple cortical cell types. While a subset of DEGs showed high expression in one cortical cell type, such as *Homer1*, which is primarily expressed in cortical excitatory neurons (Fig 2i), many other DEGs displayed a ubiquitous or mixed pattern of expression in the cortex. *Adam23*, for example, encodes an extracellular matrix protein^43^, that is expressed across many cortical clusters while *Alcam*, a cell adhesion molecule that has been found in blood-brain barrier and immune cells^44^, not only shows expected expression in non-neuronal cells, but also inhibitory neurons and several subtypes of excitatory neurons (Fig 2i). These expression patterns obfuscate the cell type origin of many CUS-associated DEGs and confound interpretation of cell type-specific CUS transcriptional networks.

### CUS-associated DEGs are cell type-specific

To date, studies measuring stress-effects on gene expression have utilized bulk tissue samples, which provide a composite of gene expression changes. We had observed from our own nuclear transcriptome profiling that bulk tissue sequencing obscured the cellular origin of CUS-induced transcriptional alterations. This prompted us to leverage our single-nucleus droplet-based RNA-sequencing (sNucDrop-seq^42^) approach in the cerebral cortex to define cell type-specific transcriptional changes that occur in adult mice exposed to CUS.

Using quality filtering settings of >600 genes detected per nucleus, we retained 31,260 neocortical nuclei (12,402 uniquely mapped reads per nucleus) from adult control (15911 nuclei) and CUS (15349 nuclei) males, detecting, on average, 2,566 transcripts per nucleus. Our analysis segregated nuclei into 26 distinct clusters, which we identified as excitatory (9 clusters), inhibitory (4 clusters), or non-neuronal (6 clusters) by their expression of known marker genes for major cortical cell types (Fig 3a,b). Excitatory neurons (*Slc17a7*+) were further sub-categorized by their superficial-to-deep layer distribution within the cortex (layer 2 to layer 6), and every major sub-class of cortical inhibitory neurons (*Gad2*+) was also captured in our analysis (Fig 3b). Non-neuronal clusters were identified as astrocytes (*Gja1*+), oligodendrocyte precursor cells (*Pdgfra*+), oligodendrocytes (Oligo1: *Opalin*+; Oligo2: *Enpp6*+), microglia (*Ctss*+), and endothelial cells (*Flt1*+). We also uncovered non-cortical contamination of our cerebral cortex dissections, such as striatal cells (~7%) and the connecting claustrum (~1%).

**Figure 3.**
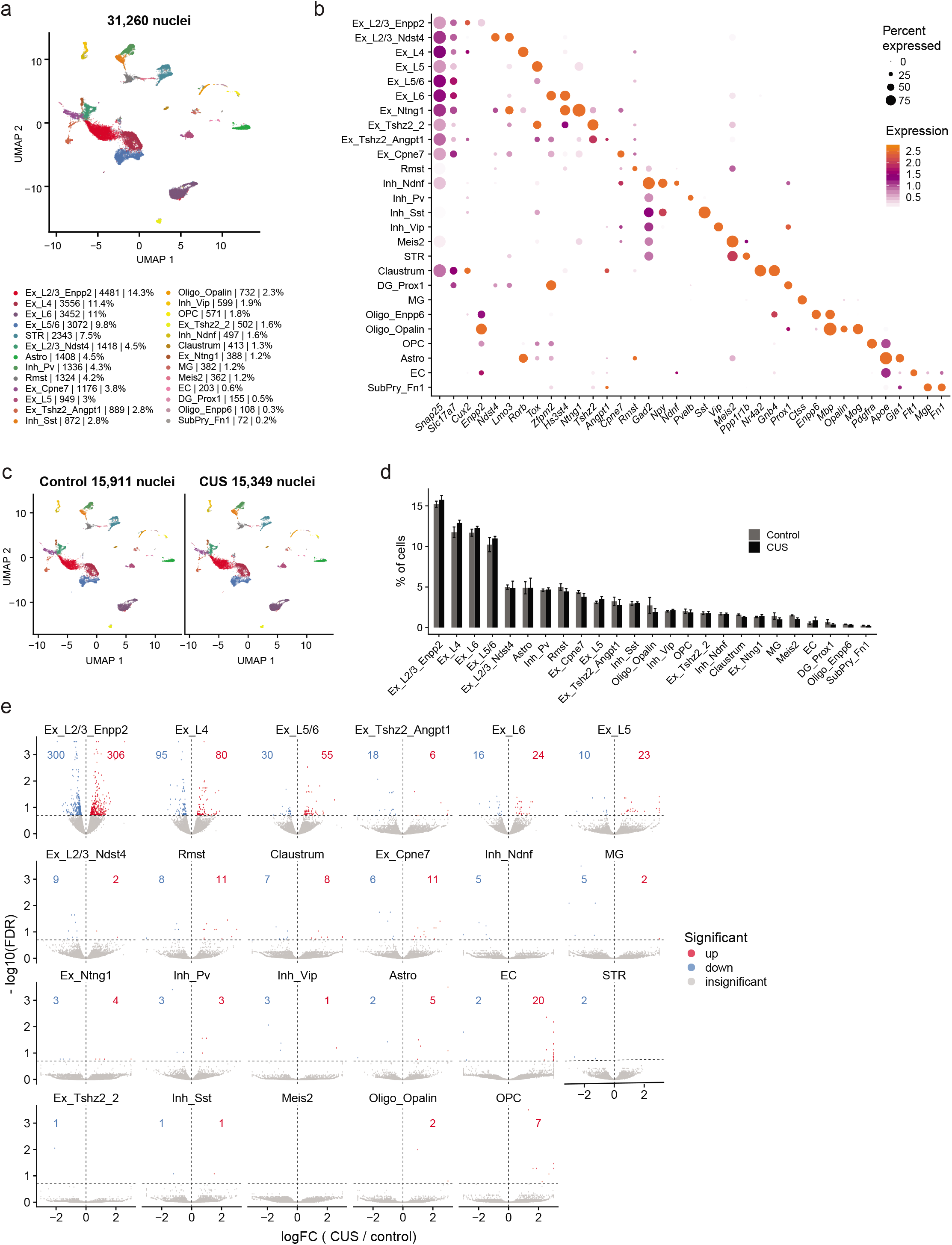
Identification of CUS-driven cortical cell type-specific gene expression changes using single-nucleus RNA-sequencing. (a) Visualization of UMAP plot displaying 26 clusters segregated from all 31,260 nuclei isolated from adult control and CUS mouse cortices (n=8 mice). Ex, excitatory neurons; Inh, inhibitory neurons; Astro, astrocytes; OPC, oligodendrocyte precursor cells; Oligo, oligodendrocytes; MG, microglia; EC, endothelial cells. (b) Bubble chart showing expression of cell type-specific marker genes for each cortical cell cluster. (c) UMAP plots depicting clusters identified from control (15,911 nuclei) and CUS nuclei (15,349 nuclei). (d) Percentage of control and CUS nuclei for every cortical cluster. (e) Volcano plots of differentially expressed genes between control and CUS cortical nuclei (FDR<0.2). Upregulated genes are shown in red and downregulated genes are displayed in blue. Error bars represent s.e.m.

Because we had previously observed discrete clustering of cortical nuclei in seizure-induced mice^42^, we asked whether we could identify CUS-dependent transcriptional states in the cortex. Upon comparing the segregation of cortical nuclei between control and CUS samples, we found that the representation and distribution of nuclei are largely similar under both conditions (Fig 3c,d), indicating that CUS exposure does not lead to altered cellular composition of the neocortex.

Classifying cortical cell types allowed us to identify cell type-specific CUS-associated DEGs by comparing the expression of nuclear transcripts between control and CUS samples for each identified cluster (Supplemental Table 3). Our analysis uncovered CUS-associated DEGs across virtually every cortical cluster. We noted that excitatory neurons displayed the most transcriptional dysregulation, as measured by DEG number, with *Enpp2*+ layer 2/3 (L2/3_Enpp2) excitatory neurons showing the greatest number of CUS-associated DEGs (n= 606 DEGs; Fig 3e). We also found a number of DEGs that are shared across discrete excitatory neuronal clusters; 36% of layer 4 DEGs (63 / 175 genes; Supplemental Fig 2a), 43.5% of layer 5/6 DEGs (37 / 85 genes; Supplemental Fig 2b), and 47.5% of layer 6 DEGs (19 / 40 genes; Supplemental Fig 2c) were shared with DEGs in L2/3. This partial overlap suggests that common molecular responses are induced across several different subclasses of cortical excitatory neurons, but that many CUS-associated transcriptional changes are unique to each neuronal subtype.

We also compared CUS-associated DEGs identified in the bulk nuclear RNA-seq and sNuc Drop-seq analyses in order to determine their cellular origin. We identified 66 individual genes that were classified as DEGs in the bulk nuclear RNA-seq dataset and also significantly dysregulated in and across several cortical excitatory, inhibitory, and non-neuronal cell populations obtained by sNucDrop-seq (Supplemental Fig 2d). A large majority of these shared DEGs showed a concordant pattern of gene deregulation (e.g., DEGs that were up-regulated in the CUS bulk nuclear RNA-seq dataset were also up-regulated by CUS in data generated by sNucDrop-seq), indicating that these DEGs are reproducibly regulated by CUS. Thus, application of these two methods of nuclear RNA-sequencing enabled discovery of transcriptional targets of CUS with cellular precision.

Considering the large number of DEGs obtained in the L2/3_Enpp2 cluster, we next examined the functional properties of these genes. We found that CUS-associated DEGs were significantly enriched in genes involved in glutamatergic neurotransmission and synapse structure, as indicated by GO enrichment analysis (Fig 4a). Notably, an enrichment of excitatory synapse-related GO terms, pathways, and gene-sets were also identified in GWAS studies of MDD^45–47^, suggesting the presence of shared transcriptional processes between CUS mice and MDD patients that impact glutamatergic synaptic function and lending further ethological validity to the CUS model of depression. Our analysis identified several DEGs previously implicated in MDD, such as *Negr1. Negr1* has been identified in multiple GWAS studies of MDD but not found to be differentially expressed in bulk RNA-seq analyses of chronically stressed mouse cortices^30^, including our own (Supplemental Table 2). Together, our data underscore the capability of single nucleus profiling in extracting subtle cell type-specific gene expression changes that are lost in whole tissue analysis.

**Figure 4.**
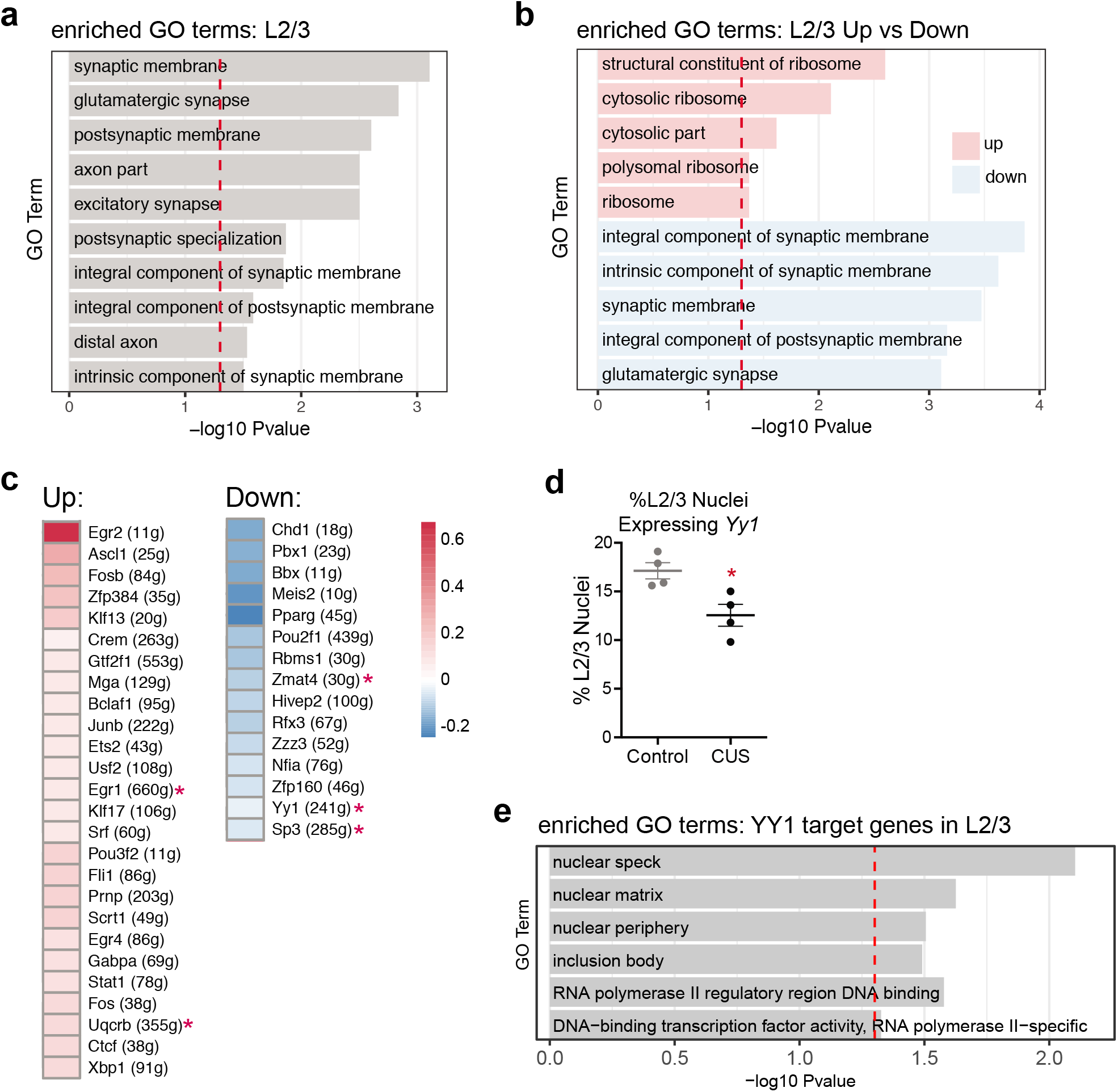
Synaptic gene expression and YY1 regulatory activity are decreased in CUS L2/3 excitatory neurons. (a) L2/3 DEGs are enriched for synapse- and axonal-related gene ontology (GO) terms (FDR<0.5). (b) Upregulated L2/3 DEGs (pink) are enriched for ribosomal-related GO terms. Downregulated L2/3 DEGs (blue) primarily encode synapse-related proteins. (c) Heatmap of CUS L2/3 SCENIC results showing 26 transcription factors that are significantly upregulated in activity and 15 that are downregulated in activity relative to controls. Legend depicts normalized mean AUC values for CUS nuclei relative to control nuclei. (d) Percentage of L2/3 control and CUS nuclei expressing *Yy1* (*P<0.05; error bars represent s.e.m.). (e) YY1 target genes in L2/3 neurons encode nuclear proteins and RNA Pol II-mediated transcription-related GO terms.

Furthermore, GO analysis on L2/3 down-regulated DEGs alone showed that the transcription of these synapse-related genes is decreased in response to CUS, in contrast to the ribosomal-related GO terms that were enriched in up-regulated DEGs (Fig 4b). The decreased synaptic gene expression in CUS excitatory neurons is consistent with previously reported observations of decreased dendritic spine density in PFC pyramidal neurons of stressed rodents and MDD subjects^27^, as well as our own data demonstrating reduced neuronal activity in the PFCs of CUS mice (Fig 2f,h). Together these data provide a potential mechanism for the known effects of chronic stress on synaptic volume and activity in cortical excitatory neurons.

### CUS decreases YY1 regulatory activity in layer 2/3 excitatory neurons

Gene expression is a precisely orchestrated and cell type-specific process—regulated, in part, by networks of transcription factors (TFs) and co-factors within the cell. To gain mechanistic insight into the deregulation of transcription observed in CUS L2/3_Enpp2 nuclei, we sought to identify specific gene regulatory networks (GRNs) that are altered by CUS in this excitatory neuronal subtype. Using SCENIC^48^, which infers enriched GRNs associated with specific transcription factors from single-cell RNA-seq data, we detected 346 TFs in L2/3_Enpp2 cortical nuclei whose binding motifs are significantly enriched in their coregulated GRNs. Among these TFs, 41 showed differential GRN activity in CUS nuclei, and 5 of these themselves were significantly dysregulated in expression in our sNucDrop-seq data (Fig 4c, Supplemental Fig 3a,b). We also identified altered regulatory activity of several TFs previously implicated in chronic stress, including FOS, FOSB, and CREM, which are significantly up-regulated in CUS L2/3_Enpp2 neurons compared to controls (Fig 4c, Supplemental Fig 3a).

Interestingly, we found that the regulatory activity of CTCF and YY1 in L2/3_Enpp2 nuclei was significantly altered by CUS in opposing directions (Fig 4c, Supplemental Fig 3a,b). This finding is consistent with CTCF and YY1’s preference for different regulatory elements; while CTCF tends to occupy insulator elements, YY1 has been shown to preferentially occupy active enhancers and promoters^49^. These proteins are of particular interest given CTCF and YY1’s role in regulating cell type-specific gene expression programs and our own data showing CTCF and YY1 motifs at regulatory regions of dynamic chromatin looping in CUS frontal cortices (Fig 2h). These findings suggest that CUS decreases YY1 binding and subsequently increases CTCF-mediated chromatin interactions at this genomic region.

Moreover, a recent study reported that a number of SNPs associated with MDD disrupt the binding motifs of these chromatin regulators^50^, indicating that perturbation of genome architecture by CTCF and YY1 may be a functional mechanism underlying MDD etiology. While the nuclear transcript levels of *Ctcf* itself were insignificantly altered between control and CUS L2/3_Enpp2 neurons, we found that *Yy1* was significantly decreased by CUS in this cell population (Supplemental Table 3). This finding was of great interest given that *Yy1* had been previously identified as a hub gene in a whole blood gene expression network analysis of MDD, and its expression was found to be negatively correlated with MDD status^51^. Further examination revealed that the decreased expression level of *Yy1* (Supplemental Fig 4a) was largely driven by a significant decrease in the percentage of L2/3_Enpp2 nuclei expressing *Yy1* transcripts in CUS conditions (Fig 4d, Supplemental Fig 4b). Furthermore, this CUS-associated decrease in *Yy1*-expressing L2/3_Enpp2 neurons captured by our single-nucleus RNA-seq was observed in both batches of sequencing that we performed, indicating that this finding is reproducible across cohorts (Supplemental Fig 4b). Together, these data demonstrate that CUS decreases YY1 regulatory activity, in part, through down-regulating *Yy1* transcription.

Despite its constitutive expression in the brain, studies of YY1 have been generally constrained to its role in early CNS development and its molecular function in the adult brain is largely unexplored. To better understand YY1’s regulatory function in adult cortical excitatory neurons, we performed a GO enrichment analysis on the 241 genes that comprise the YY1 GRN/regulon as determined by SCENIC (Supplementary Table 3). We found that YY1-regulated genes primarily encode nuclear proteins that regulate RNA Pol II-mediated transcription (Fig 4e), consistent with the enrichment of RNA Pol II-mediated transcription GO terms seen in DEGs down-regulated in our CUS bulk nuclear transcriptomic analysis (Fig 2e). These data suggest that decreased YY1 activity may, in part, regulate the transcriptional down-regulation observed in our bulk nuclear transcriptome analysis.

In accordance with YY1’s known ubiquitous expression, we detected *Yy1* transcripts in every major cortical cell type at roughly equal levels (Supplemental Fig 4c). We then compared the percentage of *Yy1*-expressing nuclei between control and CUS samples for every major subtype of excitatory and inhibitory neurons. We not only discovered that the percentage of *Yy1*-expressing nuclei decreased in CUS-isolated L2/3_Enpp2 neurons relative to controls, but that it also decreased in every CUS-associated excitatory neuronal subtype (Supplemental Fig 4d; P<0.1). Remarkably, this reduction in *Yy1* transcripts was not observed in any inhibitory neuronal subtype (Supplemental Fig 4d), suggesting an excitatory neuron-specific effect of CUS on *Yy1* expression.

Taken together, these findings indicate a potential function for YY1 in mediating the transcriptional consequences of chronic stress in cortical excitatory neurons.

### Chronic CORT administration down-regulates *Yy1* gene expression and decreases YY1 protein levels

Stress exposure alters multiple hormone signaling pathways in the brain. These include signaling mechanisms mediated by glucocorticoids, which are secreted upon activation of the hypothalamic-pituitary-adrenal axis and regulate downstream gene expression. To determine whether CUS decreases YY1 through glucocorticoids, we used primary cultures of mouse cortical neurons to assay the effect of stress-released corticosterone (CORT)—the principal rodent glucocorticoid—on YY1 in a relatively homogenous population of cells comprised mostly of excitatory neurons. In addition to YY1, we also examined the expression of *Nuclear receptor subfamily 3 group C member 1* (*Nr3c1*), which encodes the glucocorticoid receptor (GR). We used *Nr3c1* expression as a positive indicator of CORT exposure in this experimental model, given that *Nr3c1* transcription is auto-regulated by CORT-controlled feedback mechanisms^52^ and reportedly mediated by YY1 activity^53^. In agreement with these prior findings, we also found that *Nr3c1* is a gene member of the YY1 GRN/regulon (Supplemental Table 3).

We treated cortical neuron cultures with a physiologically relevant concentration of CORT (1 μM^54^) for varying lengths of time (0 hr, 3 hr, 24 hr, 72 hr, 1.5 wk) followed by real-time PCR (RT-PCR) and western blot analyses (Supplemental Fig 5a). We found that treating primary cortical neurons with 1 μM CORT for 72 hrs and 1.5 wk significantly decreased expression of *Yy1* (Supplemental Fig 5c), and *Nr3c1* (Supplemental Fig 5b) relative to vehicle treated (0 hr) cells. In contrast, 3 hr and 24 hr exposure to CORT had no effect on *Yy1* transcript levels (Supplemental Fig 5c). Importantly, we found that 1.5 weeks of CORT treatment significantly decreased YY1 steady-state protein levels as well (Supplemental Fig 5d), indicating that chronic CORT administration, but not acute exposure, decreases YY1 at both the transcript and protein levels. Together with our *in vivo* sNucDrop-seq data, these findings indicate that CUS exposure likely decreases YY1 activity in cortical excitatory neurons through prolonged production of CORT.

### *In vivo* down-regulation of YY1 in PFC excitatory neurons increases stress susceptibility in mice

The down-regulation in YY1 expression and activity that we observed in CUS cortical excitatory neurons indicated that YY1 might mediate the transcriptional and behavioral effects of stress. Thus, we next investigated the functional role of cortical YY1 *in vivo*. Due to YY1’s essential role in early cortical development and its ubiquitous expression, we adapted a genetic strategy to selectively ablate YY1 expression in PFC excitatory neurons of adult male mice. We performed bilateral PFC injections in 9-12 week old male mice carrying a floxed *Yy1* allele^55^ (*Yy1^fl/fl^*)with adeno-associated virus (AAV) expressing either *CamKII* promoter-driven eGFP (hereafter referred to as YY1-**ex**GFP) or Cre recombinase fused to eGFP (subsequently referred to as YY1-**ex**KO). This approach restricts viral expression of eGFP/eGFP-Cre to *CamKII*+ **ex**citatory neurons in the PFC (Fig 5b). AAV-infected PFC tissues from Yy1-exKO mice showed a ~50% reduction in *Yy1* gene expression compared to YY1-exGFP controls by RT-PCR (Fig 5c) as well as a ~50% decrease in YY1 protein levels (Fig 5d). These results are consistent with previous studies that have shown a ~50% reduction in CTCF, another ubiquitously expressed transcriptional regulator, in cortical and hippocampal tissues of CTCF floxed mice expressing *CamKII-Cre*^56–58^. These results demonstrate the high cellular heterogeneity of the cortex and underscore the need for studies that analyze cell type-specific functions of ubiquitously expressed factors such as YY1.

**Figure 5.**
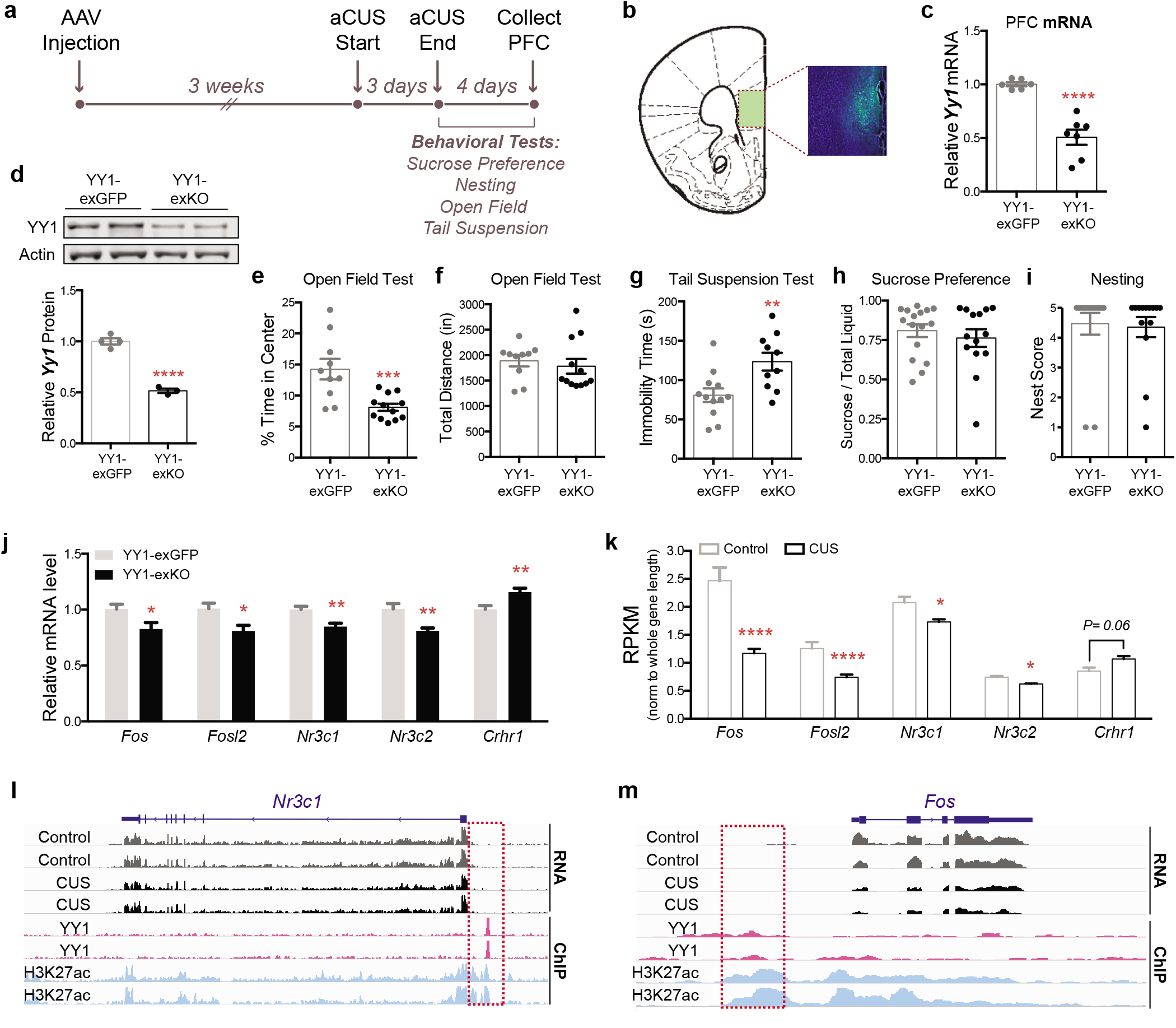
Selective genetic deletion of *Yy1* in PFC excitatory neurons enhances stress vulnerability in adult male mice. (a) Timeline of AAV injections, aCUS, and behavioral experiments. (b) Representative image of GFP expression in medial PFC of AAV-injected mouse. (c) Quantification of *Yy1* mRNA levels in YY1-exKO mice (n=7) relative to YY1-exGFP mice (n=6) (Unpaired t-test with Welch’s correction; ****P<0.0001). (d) Representative western blot showing YY1 and β-actin proteins in medial PFC tissue lysates from YY1-exGFP and YY1-exKO mice. Semi-quantification of YY1 knockdown (normalized to β-actin) is shown on the right (Unpaired t-test; ****P<0.0001; n=4 per group). (e) Decreased exploratory behavior in the open field arena in aCUS-exposed mice harboring selective loss of YY1 in PFC excitatory neurons (Unpaired t-test with Welch’s correction; P=0.001; n=10 YY1-exGFP, n=12 YY1-exKO). (f) Total locomotion in the open field test is unaltered. (g) Loss of YY1 in PFC excitatory neurons increases immobility in aCUS-exposed male mice (Unpaired t-test; P=0.007; n=12 YY1-exGFP, n=10 YY1-exKO). (h, i) Yy1-exKO males subjected to aCUS show no alterations in behavior during the (h) sucrose preference (Mann-Whitney U-test; P= 0.76; n=16 Yy1-exGFP, n=14 Yy1-exKO) and (i) nesting (Mann-Whitney U-test; P= 0.31; n=16 Yy1-exGFP, n=14 Yy1-exKO) assays. (j) Quantitative RT-PCR measurements of *Fos, Fosl2, Nr3c1, Nr3c2*, and *Crhr1* mRNA levels in aCUS YY1-exKO mice relative aCUS YY1-exGFP controls (Unpaired t-test; *P<0.05; **P<0.01; n=8). (k) RPKM values for *Fos, Fosl2, Nr3c1, Nr3c2*, and *Crhr1* PFC transcripts in control (n=4) and CUS (n=5) mice as determined by RNA-seq (*P<0.05; ****P<0.0001). (l) Snapshot of genome browser depicting nuclear RNA-seq, YY1, and H3K27ac ChIP-seq reads at the mouse *Nr3c1* locus. Overlay of YY1 and H3K27ac ChIP signal ~5kb upstream of the Nr3c1 TSS is highlighted in a red dashed line box. (m) Snapshot of genome browser depicting nuclear RNA-seq, YY1, and H3K27ac ChIP-seq reads mapped to the mouse *Fos* locus. Overlay of YY1 and H3K27ac ChIP signal ~20kb upstream of the Nr3c1 TSS is highlighted in a red dashed line box. *P<0.05; **P<0.01; ***P<0.001; ****P<0.0001. Error bars represent s.e.m.

To investigate the behavioral consequences of deleting *Yy1* in PFC excitatory neurons, we performed the sucrose preference, nesting, and open field assays in both cohorts of mice following 3 weeks of recovery (Supplemental Fig 6a). We found that *in vivo* deletion of *Yy1* in PFC excitatory neurons alone did not induce a robust depressive- and anxiety-like state in mice, as determined by the comparable sucrose preference (Supplemental Fig 6b) and open field responses (Supplemental Fig 6d,e) between YY1-KO males and YY1-exGFP controls. YY1-KO mice, however, did exhibit a trend towards decreased nesting behavior (Supplemental Fig 6c; P=0.08). These results indicate that loss of YY1 in PFC excitatory neurons alone does not significantly perturb PFC function to drive complex behaviors under these experimental conditions.

Given that our experimental and computational analyses uncovered chronic stress-associated decreases in *Yy1* transcription and regulatory activity in neocortical excitatory neurons, we reasoned that decreased YY1 function in this neuronal population might influence stress coping *in vivo*. Thus, to determine whether selective loss of YY1 in adult PFC excitatory neurons influences stress sensitivity, YY1-exGFP and YY1-exKO animals were subjected to an **a**bbreviated form of the **CUS** paradigm that consists of 3 days of CUS stressors (subsequently referred to as **aCUS**; Fig 5a, Supplemental Table 4) before undergoing the sucrose preference, nesting, open field, and tail suspension tests. Notably, we found that loss of YY1 in this cell population significantly enhanced stress susceptibility in male mice. YY1-exKO males spent less time in the center of the open field arena compared to aCUS-exposed Yy1-exGFP controls (Fig 5e) with no alteration in locomotor activity (Fig 5f). YY1 inactivation in PFC excitatory neurons also significantly increased immobility time in the tail suspension test in aCUS mice, but did not impact sucrose preference or nesting behavior compared to aCUS YY1-exGFP males (Fig 5h,i).

Having discovered that Yy1-exKO mice subjected to aCUS develop CUS-associated phenotypes, we hypothesized that their behavior was driven by a pattern of neuronal gene deregulation comparable to CUS-exposed males. First, we compared the expression of the genes encoding the stress hormone receptors, *Nr3c1* and *Nr3c2*, and corticotropin-related hormone (CRH) receptor, *Crhr1*. These genes have been extensively implicated in stress-related mood and anxiety disorders and in chronically stressed rodents. Gene expression assays performed in medial PFC tissues from Yy1-exKO males subjected to aCUS showed a significant reduction in both *Nr3c1* and *Nr3c2* and increased *Crhr1* expression relative to aCUS-stressed Yy1-exGFP controls (Fig 5j). We compared these gene expression changes to those obtained in bulk nuclear RNA-seq experiments from CUS-subjected mice. In agreement with with data obtained from aCUS Yy1-exKO mice, RNA-seq analysis of CUS PFC tissues showed a significant reduction in both *Nr3c1* and *Nr3c2* expression and a trend towards increased *Crhr1* expression (P= 0.06) compared to non-stressed control males (Fig 5k).

The stress-responsive neuronal activity genes, *Fos* and *Fosl2*, are components of the activating protein-1 (AP-1) transcriptional complex. Previously published work documenting reduced expression of *Fos* and *Fosl2* in the PFCs of chronically stressed rodents^30,59,60^ and our own data showing dysregulated FOS gene regulatory activity in CUS L2/3_Enpp2 neurons (Fig 4c) prompted us to measure *Fos* and *Fosl2* expression in aCUS-exposed Yy1-exKO animals. We found that *Fos* and *Fosl2* are both down-regulated in the PFCs of aCUS-exposed Yy1-exKO males compared to stressed Yy1-exGFP controls (Fig 5j). In agreement with this finding, PFC tissues isolated from CUS males also showed significant decreased expression of these genes relative to non-stressed controls (Fig 5k).

Given that down-regulated DEGs in L2/3_Enpp2 neurons isolated from CUS mice showed an enrichment of genes encoding synaptic proteins (Fig 4a,b), we asked if aCUS exposure in YY1-ablated PFC excitatory neurons also modified the expression of synaptic genes. We assayed the expression of the synapse-related genes, *Adam23, Grm3*, and *Nrxn1*— which encode membrane-bound cell adhesion molecules and a glutamate metabotropic receptor—and found that their expression was significantly decreased in aCUS Yy1-exKO PFC tissues compared to controls (Supplemental Fig 7a). Given that these genes are not part of the YY1 regulatory network, these data indicate that loss of YY1 can indirectly impact the expression of synapse-related genes following stress exposure. Together with the aforementioned findings, these data demonstrate that an abbreviated 3-day CUS exposure in animals harboring selective loss of YY1 in PFC excitatory neurons provokes behavioral and transcriptional responses associated with 12 days of CUS exposure.

We next sought to assess whether the dysregulated expression of genes shared between Yy1-exKO aCUS and CUS animals could be directly caused, in part, by decreased YY1 function. We surveyed the genomic landscape at the shared DEGs, *Nr3c1* and *Fos*, for enrichment of YY1, using publicly available YY1 chromatin immunoprecipitation-sequencing (ChIP-seq) data obtained from mouse cortical cells^61^. We also examined H3K27ac, a chromatin mark associated with enhancers and active promoters, in adult cortical excitatory neurons^62^. YY1 reportedly regulates the expression of *Nr3c1* in the hypothalamus^53^ and *Fos* in HeLa cells^63^, and we reasoned that YY1 also mediated the transcription of these genes in cortical cells under stress conditions. In agreement with our SCENIC analysis, which identified *Nr3c1* as a part of the YY1 GRN/regulon in L2/3_Enpp2 neurons (Supplemental Table 3), and previous findings of YY1 co-localization with H3K27ac signal, we observed enrichment of YY1 binding ~4 kb upstream of the *Nr3c1* transcription start site (TSS) that aligned with H3K27ac signal (Fig 5l). We also surveyed the genomic region surrounding *Fos* for YY1 binding. Although *Fos* had not been identified in our analysis as a component of the YY1 regulatory network, its transcription is linked to YY1 indirectly through YY1’s interactions with the ATF/CREB family of transcription factors^63^. Notably, we observed YY1 signal at an H3K27ac-enriched enhancer region ~15 kb upstream of the *Fos* TSS (Fig 5m). H3K27ac signal at this enhancer was recently linked to the activation of *Fos* expression in murine primary cultured neurons^64^, suggesting that YY1 binding at this enhancer may regulate *Fos* transcription.

### CUS drives depressive- and anxiety-like behaviors in adult female mice that are mediated by YY1 activity in PFC excitatory neurons

Given the increased prevalence of stress-related mood disorders in women, an examination of neuronal YY1 function and stress adaptation in female subjects is especially important. Accordingly, we first characterized the behavioral effects of CUS in adult female mice (9-10 weeks old) to confirm that the CUS paradigm drove the same depressive- and anxiety-associated phenotypes as was observed in male mice. Adult female mice (9-10 weeks) subjected to twelve consecutive days of CUS lost a significant amount of weight compared to age-matched controls (Fig 6b), although body weights between the two groups were evenly distributed prior to CUS (Fig 6a). Female CUS mice exhibited decreased food consumption (Fig 6c), sucrose preference (Fig 6d), and liquid consumption (Fig 6e) relative to non-stressed controls. CUS female mice also spent significantly less time exploring the center of the open field arena (Fig 6f) although they traveled a comparable distance to unstressed controls (Fig 6g). Additionally, CUS produced a modest effect on behavioral despair as measured by the tail suspension test; CUS females, like CUS males, showed a nonsignificant trend towards increased immobility compared to non-stressed controls (P=0.1; Fig 6h).

**Figure 6.**
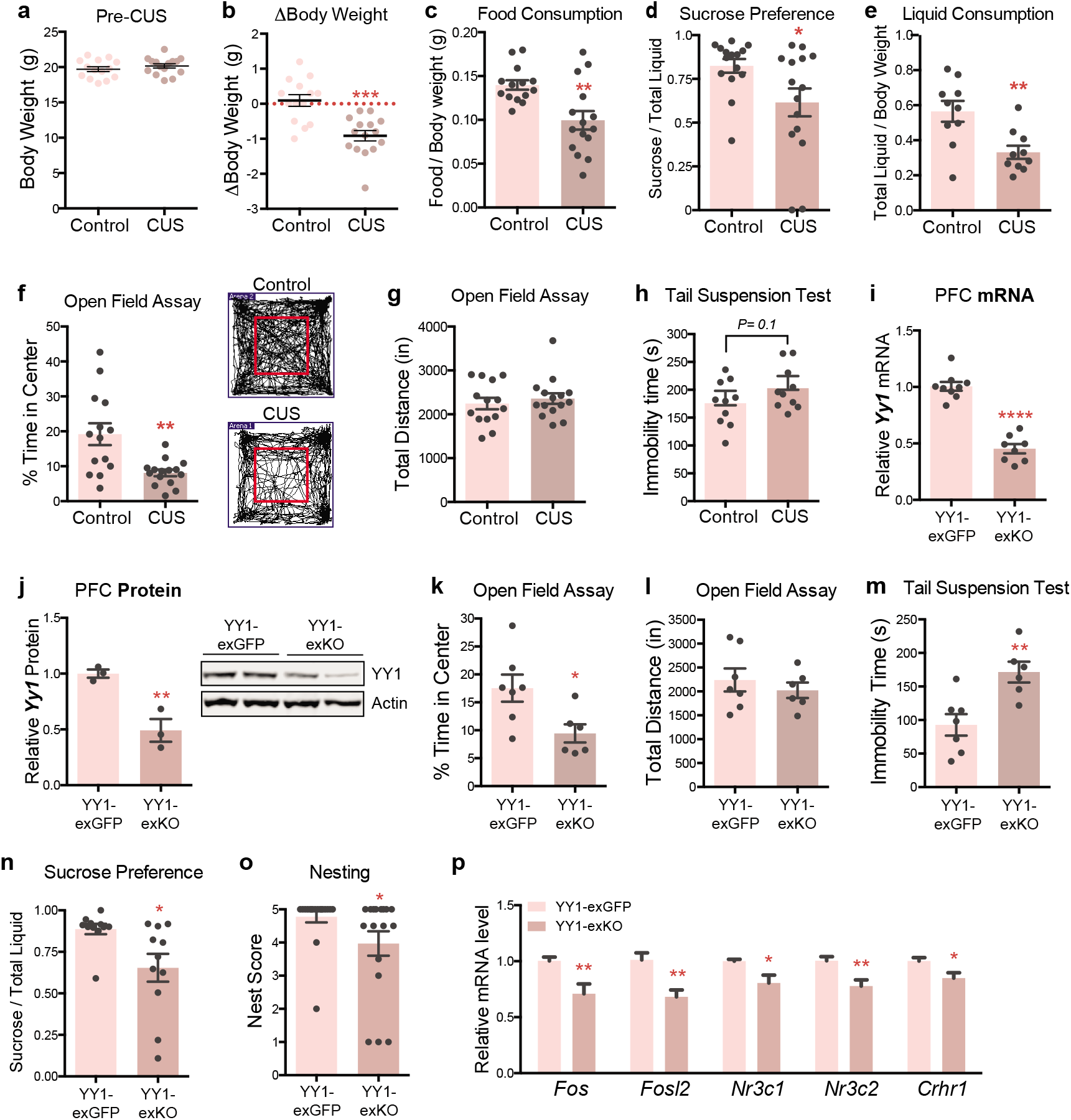
CUS induces a depressive- and anxiety-like state in adult female mice that is driven by YY1 activity in PFC excitatory neurons. (a) Pre-CUS body weights of control (n=14) and CUS (n=15) adult female mice. (b) CUS drives weight loss in adult female mice (Unpaired t-test; P<0.0001; n=14 controls, n=15 CUS). (c) Food consumption of control and CUS females normalized to body weight. (Unpaired t-test with Welch’s correction; P=0.003; n=14 controls, n=15 CUS). (d) CUS decreases sucrose preference in adult female mice compared to unstressed controls (Mann-Whitney U-test; P= 0.02; n=14 controls, n=15 CUS). (e) Liquid consumption of control and CUS females normalized to body weight (Unpaired t-test; P=0.004; n=10 per group). (f) CUS decreases exploratory behavior of adult female mice in the open field test (Unpaired t-test; P=0.004; n=14 controls, n=15 CUS). (g) Locomotor activity of female mice is unaffected by CUS (Unpaired t-test; P=0.5; n=14 controls, n=15 CUS). (h) Immobility times of control and CUS females subjected to the tail suspension test (Unpaired t-test; P=0.01; n=10 per group). (i) Quantification of *Yy1* mRNA levels in YY1-exKO mice (n=8) relative to YY1-exGFP females (n=9) (Unpaired t-test with Welch’s correction; ****P<0.0001). (j) Representative western blot of YY1 in medial PFC tissue lysates from YY1-exGFP and YY1-exKO females. Quantification of YY1 protein expression (normalized to β-actin) is shown on the right (Unpaired t-test; P<0.009; n=3 per group). (k) Decreased exploratory behavior in the open field arena exhibited by aCUS-exposed mice harboring selective loss of YY1 in PFC excitatory neurons (Unpaired t-test with Welch’s correction; P=0.02; n=7 YY1-exGFP, n=6 YY1-exKO). (l) aCUS does not alter locomotor activity of Yy1-exKO mice relative to Yy1-exGFP controls (Unpaired t-test; P=0.5; n=7 YY1-exGFP, n=6 YY1-exKO). (m) Loss of YY1 in PFC excitatory neurons increases helplessness behavior in aCUS-exposed females in the tail suspension test (Unpaired t-test; P=0.005; n=7 YY1-exGFP, n=6 YY1-exKO). (n, o) Female Yy1-eKO mice subjected to aCUS show significantly decreased (n) sucrose preference (Mann-Whitney U-test; P= 0.02; n=11 Yy1-exGFP, n=11 Yy1-exKO) and (o) decreased nesting behavior (Mann-Whitney U-test; P= 0.01; n=18 Yy1-exGFP, n=17 Yy1-exKO). (p) Quantitative RT-PCR measurements of *Fos, Fosl2, Nr3c1, Nr3c2*, and *Crhr1* mRNA levels in aCUS YY1-exKO females relative aCUS YY1-exGFP controls (Unpaired t-test; P=0.005; n=7 per group). *P<0.05; **P<0.01; ****P<0.0001. Error bars represent s.e.m.

Having ascertained that CUS drives similar depressive- and anxiety-like phenotypes in male and female mice, we next assessed whether YY1 similarly functions in female PFC cortical neurons to control stress responses. We employed the same genetic strategy that we used in males to selectively inactivate YY1 from excitatory neurons in the PFCs of female mice (Supplemental Fig 6a). PFC tissues obtained from adult female *Yy1^fl/fl^* mice infected with AAV expressing *CamKII-eGFP-Cre* (YY1-exKO) showed a ~50% reduction in *Yy1* transcript (Fig 6i) and protein levels (Fig 6j) compared to *CamKII-eGFP* infected females (YY1-exGFP). We had also observed a ~50% reduction of YY1 in AAV-infected male PFC tissues; thus YY1 levels do not significantly differ between males and females in this brain region.

We next assessed behavior in female Yy1-exGFP and Yy1-exKO mice. As we had observed in males, loss of YY1 in PFC excitatory neurons alone did not impair behavior in female animals. We observed comparable sucrose preference (Supplemental Fig 6f), nesting (Supplemental Fig 6g), exploratory behavior and locomotion in the open field test (Supplemental Fig 6h,i) between both YY1-exGFP and YY1-exKO cohorts. Remarkably, however, we found that selective deletion of YY1 in PFC excitatory neurons enhanced the stress sensitivity of adult females. YY1-exKO female mice, like their male counterparts, spent significantly less time in the center of the open field arena following exposure to aCUS than Yy1-exGFP controls (Fig 6k) without displaying altered physical activity (Fig 6l) and exhibited increased behavioral despair in the tail suspension test (Fig 6m). Notably, aCUS also drove a significant decrease in sucrose preference (Fig 6n) and nest building scores (Fig 6o), behaviors that had not been altered by aCUS in Yy1-exKO males (Fig 5h,i).

Next, we asked if aCUS exposure in YY1-exKO female mice induced a similar transcriptional pattern of stress in the PFC as it did in males. We first measured the expression of the stress-related genes, *Nr3c1, Nr3c2*, and *Crhr1*, using RT-PCR on PFC tissues microdissected from aCUS-exposed YY1-exKO and YY1-exGFP females. We found significant reduction in the expression of *Nr3c1, Nr3c2*, and *Crhr1* in Yy1-exKO female PFC samples relative to controls (Fig 6p). Moreover, expression of *Fos* and *Fosl2* were also significantly reduced in female Yy1-exKO PFC tissues (Fig 6p), as was observed in males.

Taken together, we conclude that twelve days of CUS induces stress-associated behaviors in both females and males and that these behaviors are, in part, mediated by YY1 function in PFC excitatory neurons.

## DISCUSSION

In this study, we show that twelve days of chronic unpredictable stress induces a depressive- and anxiogenic-like state in male and female mice. By performing a battery of behavioral tests and genome-wide sequencing of nuclear RNA transcripts, including the first with cellular precision, we report a novel role for YY1 in mediating CUS-induced phenotypes in cortical excitatory neurons. We also found altered patterns of chromatin interaction in frontal cortical tissues of CUS-subjected mice that were associated with neuronal inactivation, the first study to our knowledge that has implicated changes in chromatin folding in response to stress exposure. We report that cortical excitatory neurons—in particular layer 2/3 pyramidal neurons—exhibited the most transcriptional dysregulation by CUS, underscoring an enhanced sensitivity to chronic stress as well as a function for L2/3 neurons in modulating stress effects on behavior^65^. Furthermore, we found that CUS drove a significant down-regulation of synaptic genes in L2/3 cortical neurons, providing mechanistic support to previously described findings of decreased spine density and volume that have been observed in both post-mortem PFC tissues of depressed humans^66^ and stressed rodents^18,67–71^.

YY1 is a ubiquitously expressed zinc finger protein that contributes to structural enhancer-promoter interactions^49,72^, a common feature of mammalian gene control. YY1 plays a critical role in cortical development and haploinsufficiency of YY1 causes “YY1 syndrome”, a neurodevelopmental disorder characterized by intellectual disability, seizures, and behavioral impairment^73^. Despite its well-characterized role in early development, virtually nothing is known about YY1 function in the adult brain. Using single-nucleus RNA-sequencing, we found that *Yy1* expression is reduced in cortical excitatory neurons by chronic stress but not in bulk RNA-seq analysis, underscoring the cell type-specific effect of this dysregulation. Specific loss of YY1 in PFC excitatory neurons impaired stress coping ability in both male and female mice, as measured by deregulated transcription of stress-associated genes in the prefrontal cortex and increased stress-associated behaviors.

Our findings complement a previous study that identified YY1 as an upstream regulator of an MDD-associated transcriptional program in whole blood samples generated from MDD patients that also found that *Yy1* expression is negatively correlated with MDD status^51^. Together, this work indicates that decreased YY1 function is not only associated with depressive-like behaviors in PFC excitatory neurons of chronically stressed mice but may also influence MDD disease course in humans. Furthermore, most studies to date have virally manipulated gene expression *in vivo* by infecting whole brain regions to functionally validate the role of a target gene on behavior, thereby obscuring potential cell type-specific effects and contributions to the observed phenotype. In our study, we used *CamKII*-promoter driven expression of Cre recombinase to selectively delete *Yy1* from PFC excitatory neurons and demonstrate, for the first time, that YY1 function in this cell type, without manipulation of inhibitory neurons and glia, enhances stress sensitivity *in vivo*.

We show that inactivation of YY1 in PFC excitatory neurons alone did not impair behavior in male and female mice but rendered them more sensitive to stress. Although YY1 is constitutively expressed into adulthood, a recent study has shown that YY1 function is most critical during early cortical development and that neuronal dependence on YY1 decreases with age^61^. Our study demonstrates that loss of YY1 in cortical excitatory neurons does not drive functional impairment of the PFC but impairs neuronal adaptation to environmental stimuli. It also uncovered sex-discordant responses to aCUS in the sucrose and nesting assays. aCUS-subjected Yy1-exKO females showed decreased sucrose preference and nesting while the behavioral responses of YY1-exKO males in these same behavioral tests were virtually indistinguishable from Yy1-exGFP controls. These data intimate that YY1 inactivation in stress-exposed neurons affects female PFC function more broadly than in males to disrupt additional domains of behavior, also providing a potential mechanism underlying the increased incidence of stress-related mood and anxiety disorders in females. Future studies directly comparing male and female responses to aCUS, which were not performed in this study, are needed to address whether YY1-exKO females show enhanced stress-induced behavioral impairment than their male counterparts. Additionally, while our study validated YY1’s functional role in neuronal stress adaptation, it did not characterize the effects of YY1 overexpression on stress susceptibility. However, a recent study characterizing transcriptional regulation of stress resilience identified an enrichment of YY1 motifs in the promoters of stress resilience-associated genes^74^. That finding supports this study, and taken together they provide rationale for future work investigating YY1’s ability to rescue stress-induced behavioral phenotypes.

Intriguingly, our study found a sex-specific response in *Crhr1* gene expression between aCUS-exposed Yy1-exKO male and female mice. While aCUS increased *Crhr1* mRNA levels in PFC tissues of Yy1-exKO males relative to Yy1-exGFP controls, it led to decreased *Crhr1* expression in Yy1-exKO females. This dimorphic effect on *Crhr1* may be due to a sex-specific function of the CRH system in the PFC. Previous work demonstrates the presence of higher basal CRH levels in female PFCs than in males as well as sex-specific behavioral outcomes to CRH activation of neural circuits in the PFC^75^. We also observed significantly higher levels of *Crhr1* in the medial PFC of female mice compared to age-matched male littermates (Supplemental Fig 7b). This disparity in the expression and function of the CRH system in the PFC likely underlies the different effects that aCUS exerts on *Crhr1* expression in Yy1 exKO males and females.

Our study highlights the critical role of epigenetic factors in mediating cellular responses to environmental stimuli, which is an especially vital process in post-mitotic neurons. We show that YY1 activity is essential for neuronal adaptation to stress in both males and females, underscoring its generalizability as a target for therapeutic treatment. Our data also provide evidence for other factors involved in organizing chromatin structure, such as CTCF, in the stress-induced nuclear reprogramming of cortical neurons; these factors may function in concert with YY1 in re-shaping the transcriptional landscape in response to stress. Taken together, our findings show that chronic stress exerts a significant impact on PFC excitatory neurons to decrease their activity, and highlights the role of the nucleus as the dynamic center of coordinated cellular activity driving the adaptive transcriptional responses to chronic stress that ultimately modify behavior. Our study also demonstrates ethological validation of the CUS rodent model and its utility in uncovering clinically-relevant mechanistic insights into the behavioral consequences of stress.

